# Pathways to Consumers’ Minds: Using Machine Learning and Multiple EEG Metrics to Increase Preference Prediction Above and Beyond Traditional Measurements

**DOI:** 10.1101/317073

**Authors:** Adam Hakim, Shira Klorfeld, Tal Sela, Doron Friedman, Maytal Shabat-Simon, Dino J Levy

## Abstract

A basic aim of marketing research is to predict consumers’ preferences and the success of marketing campaigns in the general population. However, traditional behavioral measurements have various limitations, calling for novel measurements to improve predictive power. In this study, we use neural signals measured with electroencephalography (EEG) in order to overcome these limitations. We record the EEG signals of subjects, as they watched commercials of six food products. We introduce a novel approach in which instead of using one type of EEG measure, we combine several measures, and use state-of-the-art machine learning algorithms to predict subjects’ individual future preferences over the products and the commercials’ population success, as measured by their YouTube metrics. As a benchmark, we acquired measurements of the commercials’ effectiveness using a standard questionnaire commonly used in marketing research. We reached 68.5% accuracy in predicting between the most and least preferred items and a lower than chance RMSE score for predicting the rank order preferences of all six products. We also predicted the commercials’ population success better than chance. Most importantly, we demonstrate for the first time, that for all of our predictions, the EEG measurements increased the prediction power of the questionnaires. Our analyses methods and results show great promise for utilizing EEG measures by managers, marketing practitioners, and researchers, as a valuable tool for predicting subjects’ preferences and marketing campaigns’ success.

## Introduction

Prediction is one of the fundamental notions of scientific endeavour. It is the aspiration of many researchers and practitioners to be able to measure neural and physiological activity to predict a future decision or action of a particular person or to assess the success of possible marketing campaigns in the general population. In recent decades, extensive efforts have been invested in order to identify, using both electroencephalogram (EEG) and functional magnetic resonance imaging (fMRI), which neural factors are most crucial to the formation of subjective values that generate preferences and drive choices (Bartra, McGuire, & Kable, 2013; D. J. Levy & Glimcher, 2012). In the last decade, there is a growing attempt to use these neural factors of valuation in order to predict subjects’ future choices and marketing success at the population level (Genevsky & Knutson, 2018; Hsu & Yoon, 2015; Plassmann, Venkatraman, Huettel, & Yoon, 2015; Smidts et al., 2014). In the current study, we propose a novel strategy for analysis, and use EEG neural signals combined with machine learning algorithms to examine and identify factors and modeling approaches that would yield better and more accurate predictions of subjects’ future choices and population marketing success. Identifying the best predictive factors would help us better understand the neural mechanisms of choice, and would inform marketing and advertisement scholars in academia, and practitioners in the industry.

There are several traditional market research approaches that aim to assess market success of products. Most of these methods are behavioral, mainly marketing questionnaires, focus groups, and interviews. However, these traditional non-physiological measures are in some cases problematic and may contain various issues. For example, different preference elicitation methods can result in different responses (Buchanan & Henderson, 1992; Day, 1975; Griffin & Hauser, 1993; McDaniel & Kolari, 1987; McDaniel, Verille, & Madden, 1985), using questionnaires can be biased or inaccurate (Fisher, 1993; Johansson, Hall, Sikström, Tärning, & Lind, 2006; MacKenzie & Podsakoff, 2012; Neeley & Cronley, 2004; Nisbett & Wilson, 1977) and choices may not be incentive-compatible due to high cost or unavailability. In addition, using focus groups raises many concerns which are difficult to surmount. For instance, having one or several dominant individuals within a group, permitting only one opinion to be heard; the likelihood of group dynamics obscuring some of the more controversial perspectives, due to the tendency for participants to reproduce normative discourses; and the hefty reliance on the researcher’s subjective analysis and judgement in understanding the group interactive features (Smithson, 2000); representative focus groups can be highly difficult to assemble, particularly so since focus groups may discourage certain people from participating; and focus groups are not fully confidential or anonymous (Gibbs, 1997). In order to overcome these problems, researchers are attempting to identify neural and other physiological measurements that would predict consumers’ choices and population marketing success above and beyond the ability of traditional measures.

So far, most studies have focused mainly on using fMRI scans to obtain neural measures for prediction of preferences (Falk et al., 2015; Falk, Berkman, Whalen, & Lieberman, 2011; Falk, Morelli, Welborn, Dambacher, & Lieberman, 2013; Falk, Berkman, & Lieberman, 2012; Falk, O’Donnell, & Lieberman, 2012; Genevsky & Knutson, 2015, 2018; A. Smith, Douglas Bernheim, Camerer, & Rangel, 2014; Venkatraman et al., 2015; Webb, Glimcher, & Levy, 2013). However, an fMRI scanner has a very large fixed cost component and is restrictive in practice, severely limiting fMRI studies’ generalizability and commercial applications. It is expensive to purchase ($1M-$2M) and maintain ($100K-$150K yearly), it requires a costly dedicated facility, and it is immobile. Moreover, the cost of each subject, although varying, verge on the order of $500-$1000 per subject. Lastly, there are also technical limitations to fMRI, primarily a relatively low temporal resolution on the order of 2 seconds (Huettel, Song, & McCarthy, 2004). This resolution makes it difficult to examine the rapid dynamics of neural signals that are relevant for the neural mechanisms underlying value representation. In contrast, the most advanced EEG devices cost roughly $50K, require little support and maintenance, are portable, and have very low marginal cost for running experiments. Importantly, EEG also has a very high sampling rate (on the order of 1-2ms) which enables identification of very fast changes in the neural signal over short time scales (on the order of 50ms, Luck 2005) that may carry strong predictive information about consumer preferences and choice behavior (Luck, 2014).

Despite the fact that EEG is commonly used in the neuromarketing industry (see NMSBA Website), and that there is accumulating data linking various EEG signals with value-based choice (Dmochowski, Sajda, Dias, & Parra, 2012; Fuentemilla et al., 2013; Khushaba et al., 2013; San Martin, Appelbaum, Pearson, Huettel, & Woldorff, 2013; Sutton & Davidson, 2000), only several academic studies attempted to predict subjects’ stated preferences or actual choices (Kong, Zhao, Hu, Vecchiato, & Babiloni, 2013; Ravaja, Somervuori, & Salminen, 2013; Telpaz, Webb, & Levy, 2015; Vecchiato et al., 2011; Yadava, Kumar, Saini, Roy, & Prosad Dogra, 2017), or population marketing success (Barnett & Cerf, 2017; Boksem & Smidts, 2015; Dmochowski et al., 2014; Guixeres et al., 2017; Venkatraman et al., 2015). However, importantly, nearly all these previous studies did not examine if their prediction accuracy was above and beyond the prediction accuracy of traditional marketing measurements. To show that a neural prediction approach could be advantageous as a marketing tool, it is crucial to demonstrate that using EEG contributes to prediction well above and beyond traditional marketing techniques. The only previous study that examined whether EEG measurements could increase the predictive accuracy of population marketing success above and beyond standard non-physiological marketing measurements was unsuccessful in doing so (Venkatraman et al., 2015). Also, devising the perfect and accurate behavioral method could take an immaterial amount of trials and errors, which accurate and objective neural measures could relieve. When weighing the tradeoffs between traditional behavioral measures and novel neural measures, the industry ought to be convinced that the shift towards neural measures is indeed worth the time and effort. Therefore, one of the main aims of the current study is to demonstrate that we can use EEG signals to predict both subjects’ future choices and population marketing success above and beyond standard marketing measurements. Another novel aim of the current study is that we propose a unique approach for increasing the benefit gained from using EEG recordings for prediction. We suggest and develop a method that uses various EEG measures in order to increase the prediction power of the EEG signal. All previous studies focused their analyses on a single type of EEG neural measure extracted from the signal. We suggest that a combination of various EEG measures would increase predictive power, because each measure captures a different cognitive aspect of the valuation process that in combination generates the overall value signal that a subject construct towards a marketing message.

Hence in the current study, we construct a general set of EEG measures that is a combination of measures that were previously shown to have predictive power. Specifically, we combine information on frequency band powers for estimating valuation (Boksem & Smidts, 2015; Braeutigam, Rose, Swithenby, & Ambler, 2004; Khushaba et al., 2012, 2013; Ravaja et al., 2013; Yadava et al., 2017), hemispheric asymmetries for estimating approach/avoidance tendencies (Davidson, 1998; Laurence & Gerhold, 2016; Ravaja et al., 2013; Vecchiato et al., 2011), and inter-subject correlations (Barnett & Cerf, 2017; Dmochowski et al., 2014; Hasson, Nir, Levy, Fuhrmann, & Malach, 2004) for estimating engagement.

We propose that a combination of various cognitive aspects related to the valuation process would serve as a better predictor for both actual future choices and population marketing success.

Some of the studies that inspected EEG recordings for value information used event-related designs (Ravaja et al., 2013; Telpaz et al., 2015) while others used more ecological stimuli such as commercials (Dmochowski et al., 2014; Venkatraman et al., 2015) or movie trailers (Boksem & Smidts, 2015). Using event-related designs has the benefit of a well-controlled study, and can employ multiple repetitions to increase power (because of short trials), with very accurate timing. However, event-related designs are not as ecological as testing real world advertisements or other realistic stimuli. The predilection to use simple event-based stimuli is understandable, as real-world stimuli, such as commercials, introduce challenges for the analysis and prediction processes. These can vary from overabundant stimuli which increases perceptual responses that may conceal the valuation signal, to issues within the time domain – what are the crucial moments during the commercial that build a strong preference towards the product and brand? In this study, we chose to tackle these and other issues and use stimuli taken from real-world advertisement campaigns, such that our analyses could be implemented both in academia and industry, and our results able to provide marketers and managers with relevant insights and tools.

Moreover, all previous studies (except Guixeres et al., 2017) applied variations of standard regression techniques towards their prediction models. We propose that applying machine learning tools on these data types could generate a much more advanced, accurate, and diverse prediction models. Machine learning models, such as linear discriminant analysis (LDA), Tree-based ensembles (Boosted and Random Forest), and Support Vector classifiers (SVC) have shown to produce increasingly successful predictions in various fields (Jordan & Mitchell, 2015). We suggest that by using these techniques we will be able to identify interactions and patterns within the data that classic linear econometric and Variance Analysis (ANOVA) techniques are blind to, thus producing better prediction models with higher accuracy. Hence, another aim of the current study is to test the prediction accuracy of several machine learning approaches instead of focusing on only one standard regression method.

However, applying these machine learning models on experimental EEG data with ecological design (commercial videos), raises new analysis challenges that need to be overcome in order to generate a robust and accurate model. Not addressing these challenges would lead to inflated accuracy rates that could lead to inaccurate or wrong conclusions. For example, how to appropriately split the data to train and test sets when the data is multi-leveled: data extracted for each viewing of the same commercial (Viewing-Level), or per type of commercial (Product-Level). Moreover, how to avoid falsely enhanced prediction rates, and how to normalize EEG data within and between subjects such that the models can gain sense of the data. Through our various prediction attempts, we have identified important issues when using machine learning algorithms for prediction of EEG data, which if not accounted for, might cause biases that were not considered or accounted for in previous studies. In the current study, we highlight, address, and propose solutions to some of these challenges.

To summarize, in the current study, we focus on using machine learning algorithms to identify predictive features from various EEG measures, collected while subjects watched product commercials. We examined whether information collected while observing commercials contains features helpful in predicting future choices between the products advertised, beyond traditional measures. We inspect this both by attempting to predict subject’s actual choices after commercial viewings, and by trying to predict the products’ success in the general population, as assessed by the commercials’ YouTube metrics and results from an online out-of-sample questionnaire. Importantly, we also address various modeling challenges and examine a variety of machine learning approaches.

By applying various techniques on multiple data-types, we hope to elucidate which measurements and models are most effective and appropriate for value prediction. Potentially, this could inform management and marketing researchers as to the explanation behind predictive EEG measures and their modeling, but mostly this provides prediction tools for individual and population preferences, two efforts Yarkoni and Westfall importantly distinguish between in behavioral sciences (Yarkoni & Westfall, 2017).

## Materials & Methods

### Experimental Procedure

33 subjects (13 males) participated in the study, aged 19-41. All Subjects gave informed written consent before participating in the study, which was approved by the local ethics committee at our University.

During the first stage of the experiment, subjects watched commercials embedded in a sketch comedy series. They watched three consecutive sketches followed by three consecutive local commercials regarding food products. This cycle of sketches-commercials repeated six times. Overall, subjects watched six different commercials, three times each, for a total of eighteen commercial views and a total of twenty-seven different sketches (**Figure 1**). The length of each sketch was between 24 to 100 seconds and each commercial was between 25 to 46 seconds. The presentation order of the commercials and sketches was randomized across subjects. We designed this stage to mimic as much as possible a real experience of TV watching. The total duration of this stage was 30 minutes. While watching, subjects were connected to an 8-electrode EEG system (Neuroelectrics, Spain), at positions F7, Fp1, Fpz, Fp2, F8, Fz, Cz, Pz, sampled at 500 Hz. The recordings of the EEG from this stage were processed and used to perform predictions of subjects’ later choices and the products’ out-of-sample and population success.

**Figure 1.**
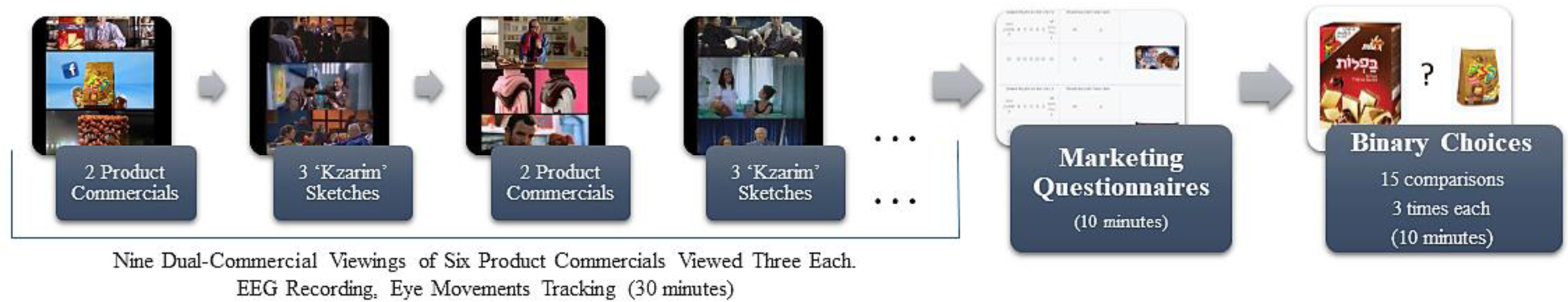
Procedure. Subjects watch six different commercials three times each, followed by a binary choice task between products that appeared in the commercials. Lastly, they filled a marketing questionnaire on each product and its commercial.

Immediately after the end of the first stage, subjects answered a standard marketing questionnaire regarding each product (see supplementary) for 15 minutes. We used the answers from the questionnaire as a baseline for our prediction attempts, which we aimed to outperform using the EEG measures we extracted while subjects viewed the commercials on the first stage. Lastly, at the third stage of the experiment, after answering the questionnaire, subjects completed a binary choice task between all product pairs. There were 6 different products, constructing a total of 15 unique product pairs. Each pair was presented 6 times, for a total of 90 binary choice trials. At the end of the experiment, one binary choice trial was chosen randomly, and the subject was given the product she chose on that trial. We used subjects’ choices from the third stage to obtain the rank order of product preferences for each subject. We then attempted to predict this subject-specific ranked order preferences using the EEG data collected while subjects viewed the commercials on the first stage.

### Out-of-Sample Metrics

The same marketing questionnaire was also completed by an online cohort of 172 subjects, to obtain an out-of-sample preference ranking regarding the products. Moreover, we gathered metrics (see below) from each commercial’s YouTube video, which served as a proxy for how popular and favorable the commercials and products are in the population.

### EEG Preprocessing

All EEG recordings were referenced to the Cz electrode and underwent 0.1 Hz high-pass filtering and a 50 Hz notch filter. Then, signals were transformed through Independent Component Analysis (ICA), and the component most related to eye movements and blinks was removed. Next, we performed Raw Data Inspection to mark apparent artifacts for removal in later processing. Finally, MATLAB’s “spectrogram” function was applied to perform Short-Time Fourier Transform (STFT), on each electrode separately, with a window of 2 seconds (1000 samples), and maximal overlap (999 samples). As the function outputs the signals’ power in various frequencies, power signals were then aggregated into well-known EEG frequency bandwidths (Luck, 2005), by choosing the signal of the frequency with maximal activity in each bandwidth. The ranges of the bandwidths we chose were: Delta 1-3.5 Hz; Theta 4-7.5 Hz; Alpha 8-12 Hz; Beta 13-25 Hz; Gamma 26-40 Hz. The final product of the preprocessing stages was power signals in the five frequency bands for each electrode and each commercial viewing separately, for every subject.

### Features: EEG Measures

We extracted the main measures from the EEG signal that previous literature has identified, to maximize our prediction accuracy.

1. **Frontal Band Powers (FBP) -** Powers for each frequency band (Delta, Theta, Alpha, Beta, and Gamma) were averaged across time per commercial viewing (Boksem & Smidts, 2015; Braeutigam et al., 2004; Khushaba et al., 2012, 2013; Koelstra et al., 2012; Luck, 2014; Ravaja et al., 2013; M. E. Smith & Gevins, 2004; Telpaz et al., 2015; Yadava et al., 2017). In order to increase our external validity, we focused on the frontal electrodes to resemble various EEG systems with only a few electrodes that are commonly used in the industry. That is, we extracted EEG data from the 3 frontal electrodes – FP1, FP2, FPz, yielding a total of 15 (5 bands X 3 electrodes) features per commercial viewing.
2. **Hemispheric Asymmetry -** We calculated the difference between band powers of the most fronto-lateral electrodes in our setup, F7 and F8, for each of the power bands, yielding 5 additional features (Cartocci et al., 2016; Davidson, 1998; Koelstra et al., 2012; Ohme, Reykowska, Wiener, & Choromanska, 2009, 2010; Ravaja et al., 2013; Sutton & Davidson, 2000; Vecchiato et al., 2010, 2011; Venkatraman et al., 2015).
3. **Inter-subject Correlations (ISC) -** For each subject, we took their power time series for a particular product commercial viewing, and cross-correlated this time series with the averaged power time-series of the same commercial viewing from all of our subject pool. For each cross-correlation calculated, we excluded from the group average the subject whose power time-series we cross-correlated with. From the resulting cross-correlation, we took the maximal correlation within 2 seconds before and after zero lag, for each frequency band, yielding a total of 5 ISC scores per commercial viewing for each subject (Barnett & Cerf, 2017; Dmochowski et al., 2014; Hasson et al., 2004).

### Features: Questionnaire (Quest)

Questionnaire responses were aggregated into eight general measures which the questionnaire was comprised of: Commercial Negative Elicitation, Commercial Positive Elicitation, Product Liking, Purchase Intent, Product Recall, Familiarity, Commercial Overall impression and Product Overall Impression (See supplementary for the full questionnaire). Afterwards, we preformed Principal Component Analysis (PCA) on all general measures from all subjects, and extracted scores for the first three components, which together exceeded 75% of explained variance. The components’ scores were used to perform predictions.

### Labels: Subject Rankings

Based on each subject’s choices in the binary choice task in the third stage, we ranked ordered each subject’s preferences for the six different products that appeared in the commercials. We ranked the responses from the binary choice task for each subject, such that the product chosen the most out of all comparisons was ranked first (Rank 1), and the product chosen the least was ranked sixth (Rank 6). This yielded six rankings per subjects, which we used as labels for predictions.

### Labels: Out-of-Sample Rankings

We aggregated the responses from the online cohort of subjects that answered the marketing questionnaire into overall rankings of the six products. We then used the aggregate overall rankings (between 1-6) as out-of-sample preferences between products. We used these overall rankings as labels for prediction.

### Labels: YouTube metrics

We collected metrics from the YouTube page of each commercial serving as a proxy for how favorable and popular each commercial was in the population. The metrics we collected were: number of likes, dislikes, shares, comments and views (views were weighted down the longer the commercial has been online). For each commercial, we divided each metric by the sum of the metric for all commercials, to gain its relative metric. Then, we performed PCA on the relative metrics, and used the summed score of the two first components as final aggregate scores. These aggregate scores were finally converted into six ordered rankings of the products’ YouTube success. We use these overall rankings as labels for prediction.

We attempt to predict all three label types, using all the features extracted from our experiment.

### Prediction Models

We attempted various machine-learning techniques, in search for the model that best suited our data and achieved highest accuracy in predicting the labels we defined, based on various feature combinations. Note that some models perform a binary classification, meaning classifying data into two possible classes (labels), some perform a multi-class classification, classifying data into more than two classes, and other perform an ordinal classification, classifying data into multiple classes which have meaningful order between them. These included the following models:

### Binary classification (prediction of two classes) models

Support Vector Machine (SVM) with a linear kernel, Logistic Regression (LOG), Boosted Decision Trees with Adaboost M1 (TREE), with 100 trees and a minimum leaf size of 5 for regularization, and K-nearest Neighbor (K-NN) with K=5.

### Nominal multi-class classification (prediction of independent categories) models

one versus all support vector machines (SVC1VA), Multinomial Regression (MNR), K-NN, and Boosted Decision Trees with Adaboost M2 (TREE) and 100 trees.

*Ordinal classification models:* These models consider the ordinal nature of rankings, our prediction classes, such as ordinal support vector machines (SVOREX), and Kernel Discriminant analysis for ordinal regression (KDLOR), with radial basis function as kernel.

Most of these models’ description and code may be found online thanks to the AYRNA research group (Gutierrez, Perez-Ortiz, Sanchez-Monedero, Fernandez-Navarro, & Hervas-Martinez, 2016) and the rest were obtained with the relevant MATLAB functions. We used the default values for all unspecified parameters.

### Modeling Approach

The final data matrix included several types of features, and 549 rows for predictions on all six ranks (31 subject’s X 6 Product Commercials X 3 viewings per commercial = 558 rows; 9 rows excluded due to extreme noise in EEG recordings), and 183 rows for binary predictions between two ranks (31 subject’s X 2 Product Commercials X 3 viewings per commercial = 186 rows; 3 rows excluded). Two subjects out of the 33 were excluded from analysis due to lost recordings. Thus, each row constituted a sample containing features from a specific product viewing of a particular subject and was assigned a label according to the rank given to that product by the subject, or by the aggregate population metrics. The features per sample include the questionnaire responses to the product viewed, and all EEG measures extracted from product commercial viewings. We performed the predictions by splitting the data randomly into ~15% test set and ~85% train set, while taking care to adhere to principles raised in the “modeling challenges” section. That is, for each prediction we randomly excluded to the test set all three viewings of one random product commercial per subject, for 6 randomly chosen subjects in binary predictions (16.4% of 183 rows of data to test) and for every subject in multi-class predictions (16.7% of 549 rows of data to test).

We trained the models on the train set, used it to predict the test set, and compared predictions to the test set labels to obtain prediction accuracy – the percentage of correct predictions, or RMSE – the root mean square error. Our model training and predictions were performed on various combinations of features and tested for significance via permutation testing (Good, 2013; Nichols & Holmes, 2002). For each feature combination, we repeated the random train-test split and prediction 10000 times, to form a distribution of accuracies that was unbiased to any specific train-test split. The accuracies presented are the distribution means and standard errors. Also, we performed the exact same procedure after shuffling all labels of the data. This enabled us to obtain a randomized ‘baseline’ distribution of predictions to improve upon with real unshuffled predictions, that provides an empirical approach to deriving statistical significance (Golland & Fischl, 2003; Ojala & Garriga, 2010), particularly for relatively small datasets (Combrisson & Jerbi, 2015). Thus, p-values were calculated as the proportion of random predictions that were higher than the mean of real predictions.

### Modeling Challenges

1. *Avoiding leakage on Multi-Leveled Experimental Data.* Since each product commercial was viewed three times, we treated the features extracted for each viewing as a separate sample (each row in **Figure 2.A**, for the 4 left columns). However, questionnaire responses are per product commercial and do not vary across the three repeated viewings of the same commercial. Therefore, we had to replicate the questionnaire responses, such that for a particular subject the same questionnaire features appeared in all three samples of viewings of the same product (Questionnaire column in **Figure 2.A**). This is a Multi-Leveled data problem – for each subject, the EEG measures are per unique product commercial viewing, but the questionnaire responses are per product commercial. While standard regression analyses can handle this with ease by clustering the errors per commercial and per subject (i.e. building a panel data), this problem is not trivial when using machine learning models, and it must be handled carefully. For example, when we randomly split the data into train and test datasets while each individual sample has the same probability to appear in either datasets, we were able to achieve **100% accuracy** in all prediction attempts that included the questionnaire data. This occurred because on some occasions, one sample out of the three viewings of a particular product by a subject was taken to the test set (indicated as the test sample in **Figure 2.A**), while the other two remained in the training set, but all three contained the same replicated questionnaire responses. The prediction model could easily employ these replicated responses to achieve high accuracy in prediction, without actually learning or predicting anything. By doing so, the model can discard all other features, since it has perfect knowledge to predict from train to test (the replicated responses). This resulted in highly inflated accuracy (using row 1 and 3 questionnaire data to predict on row 2, in **Figure 2.A**). Splitting multi-leveled data without care, can easily cause data leakage from the test set to the train set, yielding inflated false results. To address this issue, we split the data to train and test by randomly choosing sets of three samples of a given commercial that contained measurements from the three viewings of the same product commercial of a given subject, and the replicated questionnaire measurements. By doing this, we prevent any replicated data to exist both in the train and test sets, and hence prevent leakage entirely. This means our predictions were based on training sets that did not hold any information on a product commercial viewing of the subject being predicted in the test set. In essence, prediction for a specific test sample – a subject’s product commercial viewing – was based on his data from viewing other product commercials, and data from other subjects viewing the same product commercial (and others). This inherently made predictions more difficult but avoided falsely enhanced accuracy.
2. *Between Subject Normalization.* Standard regression techniques allow different intercepts and slopes for each subject to account for between subject variability that would otherwise hinder an all-inclusive general regression model. However, there are limited solutions to handle this issue when applying machine learning models on experimental data. In our experiment, each subject’s mean band powers may have a different baseline activity, may vary differently, or exist in disparate ranges. For example, the mean result in EEG power for one subject’s most (Rank 1) and least (Rank 6) preferred products could be within the range of 0-3 log-powers, while for a different subject, the least and most favorite products may be in the range of 4-10 log-powers. When the ranges of EEG powers greatly vary between subjects, it limits the machine learning algorithms’ ability to find optimal separation of the data space (**Figure 2.B, top axis).** While individual powers vary in their range and variance, they sustain their ordinality. Therefore, in order to create effective hyperplanes that separate rankings for a varied pool of subjects with different ranges, we performed a transformation which retains the powers’ ordinality while converting each subject’s powers into a common scale between 1 to 6 (**Figure 2.B, bottom axis**). We transformed the powers to a common scale, per viewing repetition separately in order to avoid averaging between viewings and maintaining data per sample. That is, for each subject, we took the powers of the first viewing (out of the three viewings) of all product commercials and rank-ordered them from 1 to 6. Thus, the first viewing of a product commercial that had the highest log-power, received the value 1, and the first viewing of a product commercial that had the lowest log-power, received the value 6. Next, we took the powers of all the second viewings of each product commercial per subject and ranked them as well from 1 to 6. We preformed the same for the third viewings of each commercial. Hence, we created a new measure, the *power-order* of each commercial viewing. Importantly, this procedure shifted all power ranges of different subjects to a common scale. Moreover, by converting powers to ranks for first, second, and third viewings of product commercials separately, the conversion cleared any habituation effect that might appear between viewing of the same product commercial. This way, band powers recorded from a first and novel viewing of a commercial were not compared against a second, more habituated, viewing of a different product commercial, but rather only against the first viewings of all other commercials. The same effect applies for the second and third viewings.
3. *Within Subject Normalization.* A machine learning model will be oblivious to the fact that three specific viewings are of the same product commercial and by the same subject. This information may be represented in the data by adding logical indicator vectors, or conversely by representing this information in some other way prior to learning a model. Since adding logical vectors is costly and increases the dimensionality of the data (in our case, 30*6 additional dimensions), we introduce a simple method that incorporates this information into the model meaningfully, without any additional dimensions. The method moves all *power-orders* of the three viewings of a certain product commercial closer to their mean across viewings, for a given subject, by half of their distance to that mean (**Figure 2.C**). Centering the *power-order* has the effect of transferring information between viewings of the same product by the same subject, similar to the effect of adding logical indicator vectors, thus taking advantage of the fact that commercials were viewed repeatedly by the same subject when the data enters our machine learning model. Thus, after we converted all EEG measures to *power-orders* as described in *Modeling Challenge 2*, we centered the *power-order* as follows:

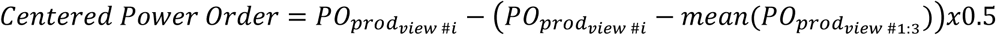 Where PO signifies *power-order*, the order-transformed band power of a specific commercial product (prod) on a certain viewing of that commercial (view #i). A product’s *power-order* was no longer just the order, but a centered order that was slightly pushed towards the *power-order* obtained for that product in other viewings of it by the same subject.
4. *Avoiding Peeking when Transforming or Selecting Data*. In our type of data, performing predictions on a subset of the data that was chosen according to its labels before prediction, and also performing transformations on that chosen subset alone, is considered a case of “Peeking”. Selecting a subset of the data by its labels before prediction gives our model a false advantage in prediction, since doing so informs the model that the data belongs to our chosen labels, and not all possible labels. In this study, when we included only a particular subset, such as viewings of products ranked highest and lowest (ranks 1 and 6), then we encountered the problem described above when predicting on that subset. Thus, in order to avoid this problem, we only interpreted the results of predictions on these subsets comparatively, against predictions on other subsets which had the same advantage (such as ranks 3 and 4).

Moreover, any data-relative transformation performed on that subset of data (as described in *Modeling Challenge 2 & 3*) will also falsely benefit the model, as these transformations will ease the separation between data samples from the chosen labels. They artificially infuse the subset of data with information on the labels, causing further ‘peeking’ which inflates accuracy when predicting on that subset. Indeed, as a demonstration of this problem, results in **Figure 2.D** show that once transformations were done after subset selecting, accuracies were significantly higher, showing the effect of further ‘peeking’ into labels.

**Figure 2.**
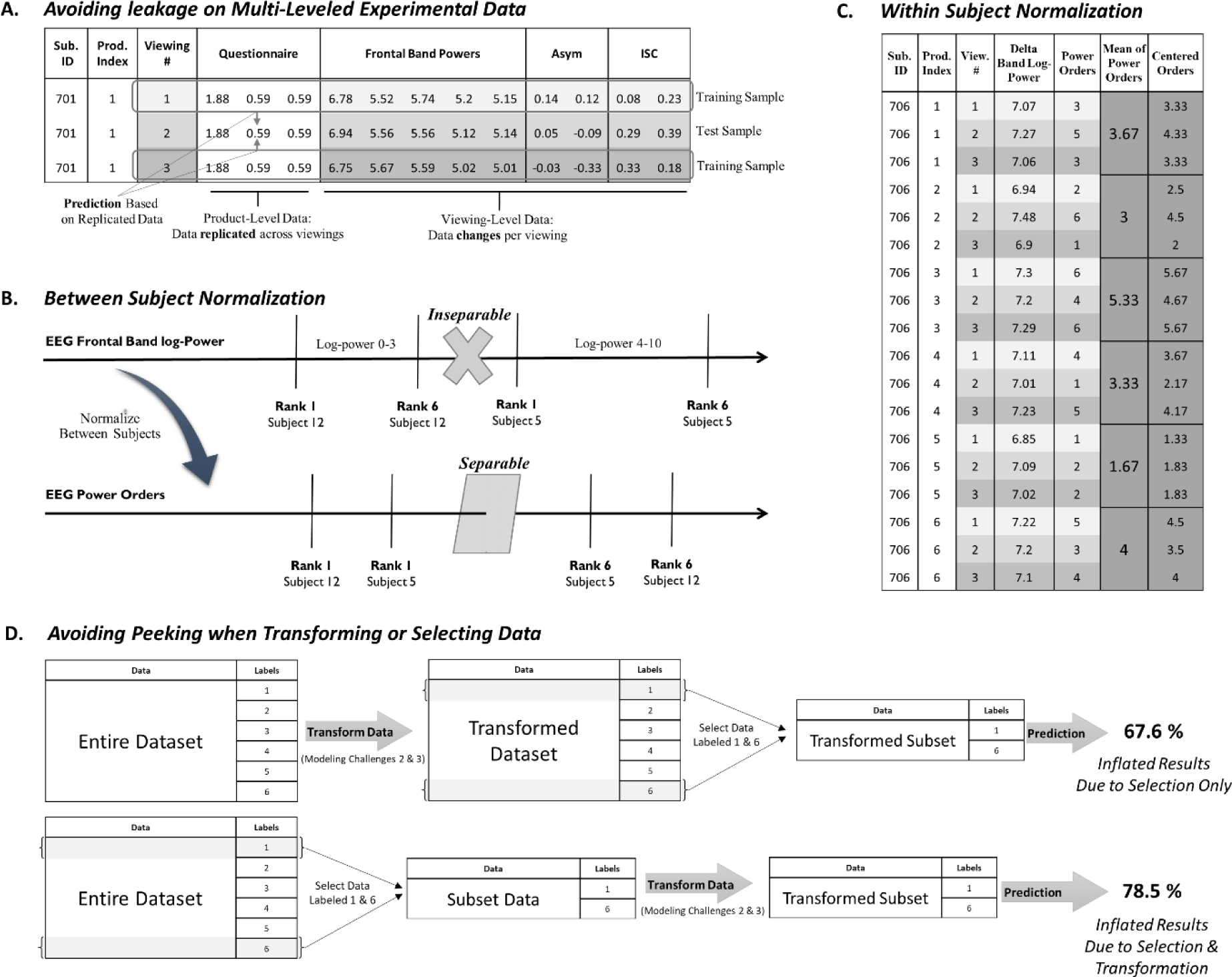
Modelling Challenges. **(A)** Every viewing of a particular product had different EEG measures but replicated Questionnaire responses – creating multi-leveled data. If we were to separate data of one viewing of a product into the test set, while leaving the other two viewings in the train set, prediction would be only based on replicated data, and therefore meaningless. **(B)** Applying machine learning models required normalization between subjects’ personal EEG responsivity patterns, otherwise separating EEG measures by their ranks would be impossible. **(C)** Normalization within subjects is required to incorporate the information that recordings came from viewings of the same product by the same subject. This panel illustrates the normalization procedure for the Delta Band log-powers of a certain subject. Within a subject, powers of the same viewing number were ordered between all different products, to create the measure “Power-Orders”. Then, power-orders of the same product, still within a subject, were centered towards their mean. **(D)** Selecting a subset of data that is labeled 1 and 6 causes inflation in accuracies, while transforming the data *after* selection causes further inflation.

## Results

### Within-Subject Pool Preferences

For meaningful predictions to be possible, the correlation of product preferences across subjects must be relatively low, such that each subject ranks the products by her own idiosyncratic subjective evaluation. It is problematic that all subjects have similar preference ordering of the products. That is, if all subjects rank the same product as the most preferred, then predicting that product’s individual ranking from EEG measures could be simply a matter of identifying its commercial’s particular pattern of neural activation, rather than basing prediction on the value signal it elicited. **Figure 3** shows the mean ranking of each product, averaged over all subjects’ rankings. Five out of six products are well within each-other’s confidence intervals, while only one product (“Candy”) appears to be significantly better than all other products. This means that for the most part, there’s high variation of product preferences between subjects, such that most products on average are liked the same. This variation in preferences between subjects ensures that we are modeling and predicting their subjective values rather than specific attributes of the product’s commercial that subjects’ EEG recording might show response to.

**Figure 3.**
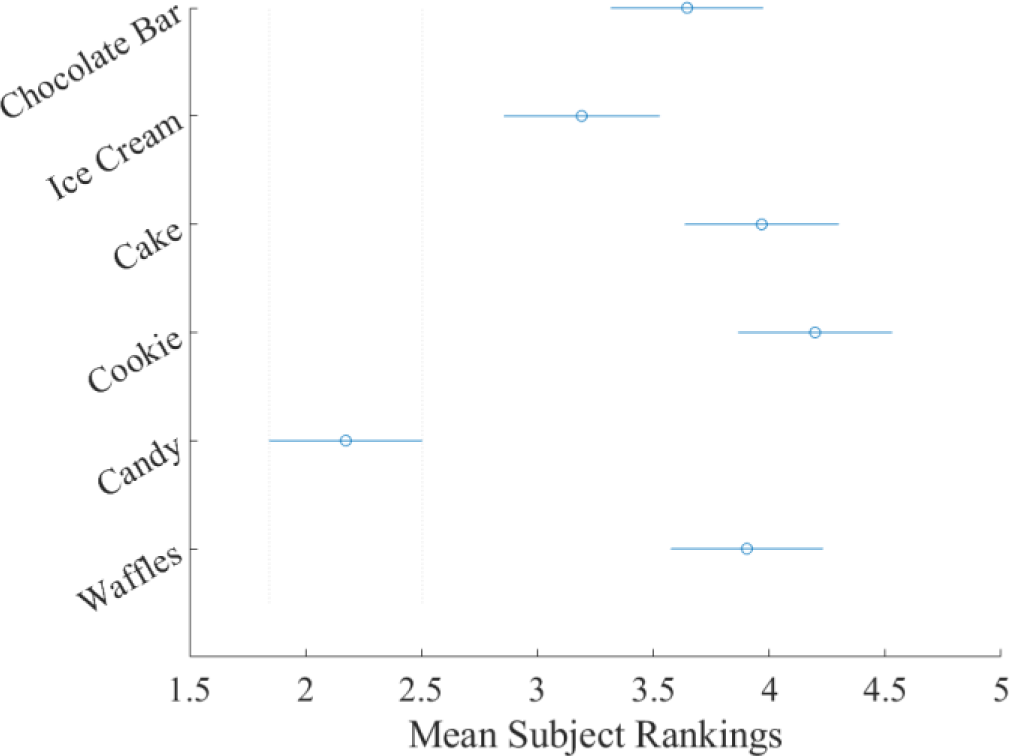
Mean Rankings per Product, Across Subjects. Mean rankings and confidence intervals (alpha = 0.05) for each product in our experiment. Only “Candy” was significantly different than all other products, reaching the best (lowest) ranking of 2.17. (F(30,5) = 20, p<0.001 on repeated measures ANOVA).

#### Questionnaire

Before attempting prediction with all our measures, we explored the relationship between components we extracted from the questionnaire and EEG with subjects’ product rankings, as obtained from the binary choice task. It is common practice to explore the relationships between the data and labels, before forming prediction models, to enhance, select and properly engineer our features. PCA components of subjects’ questionnaire responses were only moderately correlated with subjects’ product rankings as obtained from their choices, with a correlation coefficient of ρ=0.3 (Repeated Measures Correlation, p < 0.01, (Bakdash & Marusich, 2017) for the first component, and a ρ=-0.1 for the second component (n.s.) (**Figure 4**). This shows that the scores of the questionnaire are indeed related to subjects’ actual choices and can be valuable predictors of preferences. Yet, the relatively moderate correlation hints that there is enough room to improve upon using predictive measures of a different type.

**Figure 4.**
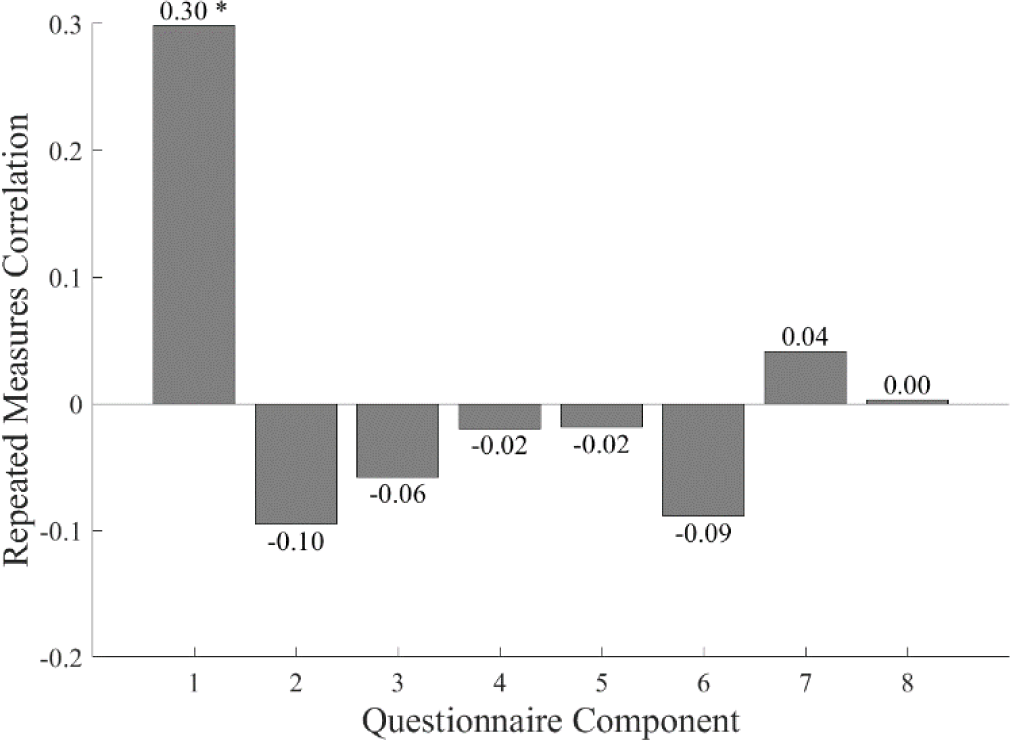
Relationship between Subject Product Rankings and Questionnaire Principal Components. The figure shows Repeated Measures Correlation coefficients (Bakdash & Marusich, 2017) between product rankings, as obtained by actual choices from the binary choice task, and the Questionnaire’s PCA component scores. A moderate connection was found, leaving room for improvement in prediction using neural measures (* = p < 0. 01). Only the first 3 components were used in predictions.

### EEG Measures

We examined our various EEG measures for relationships to subject product rankings. Previous studies have shown (I. Levy, Lazzaro, Rutledge, & Glimcher, 2011; Telpaz et al., 2015) that as the distance between preferences increase, so does the “neural distance”, as measured by neural activity, and therefore prediction accuracy increases. As such, we would expect that for product commercials ranked highest and lowest, our EEG measures would be most disparate, while for those in medium ranks, very little difference would emerge. Indeed, for Delta, Theta, and Alpha frequency bands we found a significant difference (t(31) = 2.73; t(31) = 2.77; t (31) = 3.18 respectively, p < 0.05; FDR corrected (Benjamini & Hochberg, 1995)) in EEG activity when watching commercials for distinct and distant product rankings (1 and 6), but not for close and mid-ranged rankings (3 and 4, p = n.s) (**Figure 5.A**). Furthermore, our ISC measure showed a significant difference for the distant rankings in the Gamma band (t(31) = -2.76, p < 0.05; FDR corrected), and a marginally significant difference for the Alpha band (t (31) = -1.82 p = 0.09; FDR corrected) **(Figure 5.B).** Lastly, there was a significant difference in the hemispheric asymmetry measure in the Delta and Theta power bands (t(31) = -2.55, t(31) = -2.35, p < 0.05; FDR corrected), and a marginally significant difference for the Gamma band (t(31) = 2.08 p = 0.07; FDR corrected), again in distant rankings, but not in the mid-range ranks **(Figure 5.C)**. This demonstrates the potential for some measures of the EEG recording to assist in predicting choices, but mainly for well distinct preferences, as has been shown in previous studies (I. Levy et al., 2011; Telpaz et al., 2015). Moreover, since each neural measure had at least one significant difference between rank 1 and rank 6 for a power band, this demonstrates the possible advantage of using multiple EEG measures for prediction.

**Figure 5.**
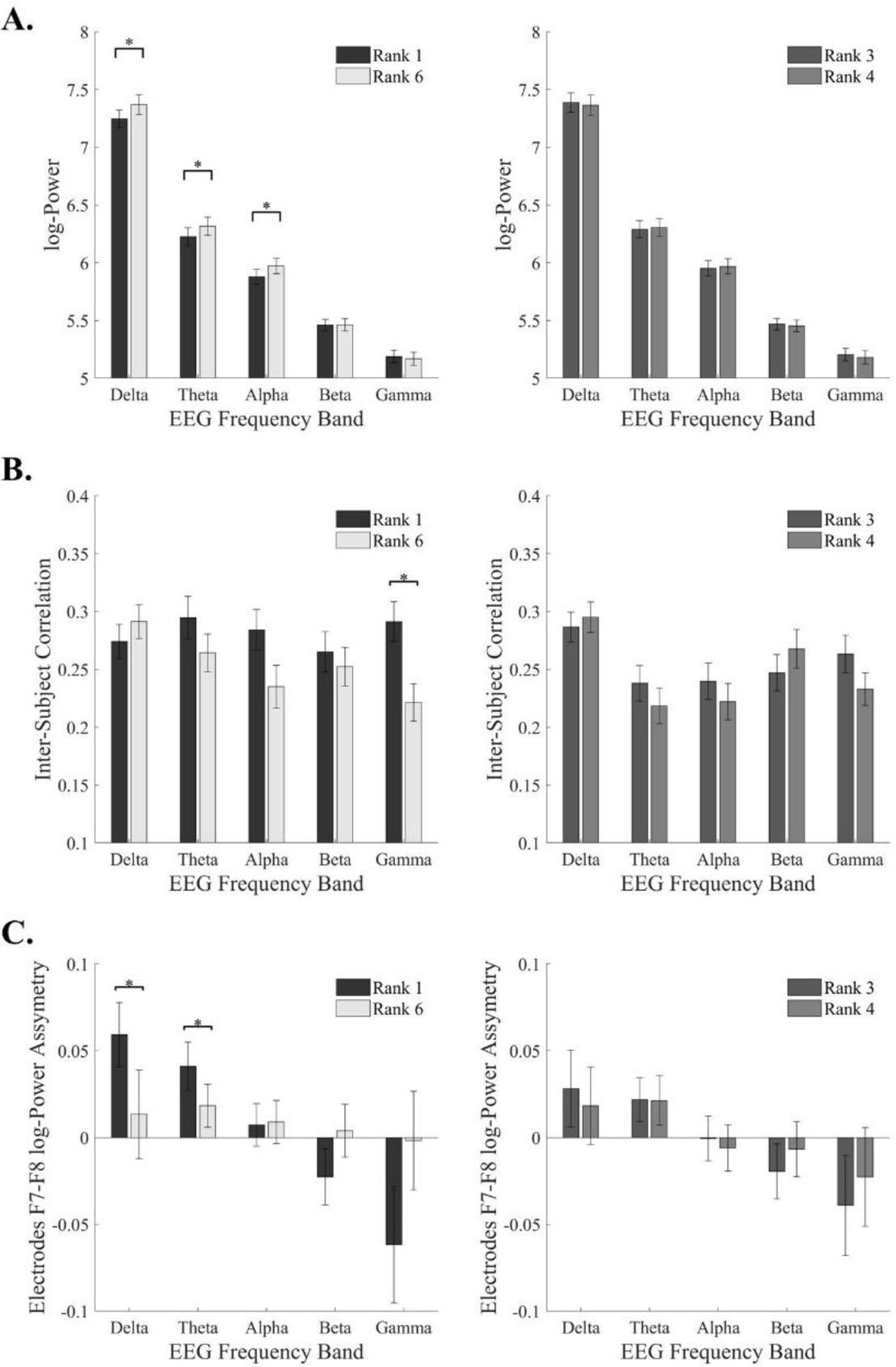
EEG Measures Differences for Distant Ranks and Mid-Adjacent Ranks. **(A)** When viewing product commercials which subjects ranked as most favorite (Rank 1), their EEG activity was significantly lower in bands Delta, Theta and Alpha, compared to products ranked least favorite (Rank 6)

This result did not replicate for Ranks 3 and 4. **(B)** Inter-Subject correlations were significantly higher for Rank 1 than Rank 6 in Gamma band, and marginally in the Alpha band (p = 0.08). No significant results for rank 3 and 4 comparisons. (**C**) Asymmetry was significantly larger only for rank 1 and 6 comparisons in Delta and Theta bands. * = p < 0.05, two-tailed paired t-test, Corrected for multiple comparisons by Benjamini-Hochberg False Discovery Rate (Benjamini & Hochberg, 1995). Error bars indicate Standard Error of Mean.

### Prediction Strategy

Our main strategy for prediction using the different machine learning models was first to examine the prediction accuracy of only the questionnaire measures in order to acquire a baseline/benchmark for prediction using a standard marketing tool. Second, to examine the prediction accuracy of the combination of the different EEG measure types (FBP, ISC, Asymmetry) on their own. This was to test if the EEG measures by themselves have predictive power. Finally, we combined both the EEG and questionnaire measures and tested if the prediction accuracy increased compared to when using the questionnaire measures alone. This is the most important comparison throughout all of our analyses because the main aim of the current study is to show that we could predict above and beyond standard marketing questionnaires. Note, that all reported prediction results are after we accounted for the modeling challenges raised in the methods sections.

### Within-Subject Prediction – Binary Classes

We applied our binary machine-learning models (SVM, LOG, KNN, TREE) on different combinations of the data types in order to predict between two product rankings at each combination, for all subjects (**Figure 6**). When trying to predict between rank 1 and 6, the combination of the EEG measures (FBP, ISC, Asymmetry) were successful in prediction on their own, reaching up to 66.27% accuracy at best, while the questionnaire measures reached 64.42% accuracy at best. Importantly, the combination of both EEG and Questionnaire measures yielded the best result, leading to 68.51% accuracy. This showed, as anticipated, that our neural measures did contribute to prediction, and therefore contained value information beyond what is captured by traditional measures alone. Moreover, our models yielded higher accuracy predicting from EEG Measures alone than from the questionnaire alone. This suggests that utilizing measures that capture various characteristics of the EEG signal is advantageous for prediction, in contrast to previous research which focused on only one EEG measurement type for explanatory purposes.

**Figure 6.**
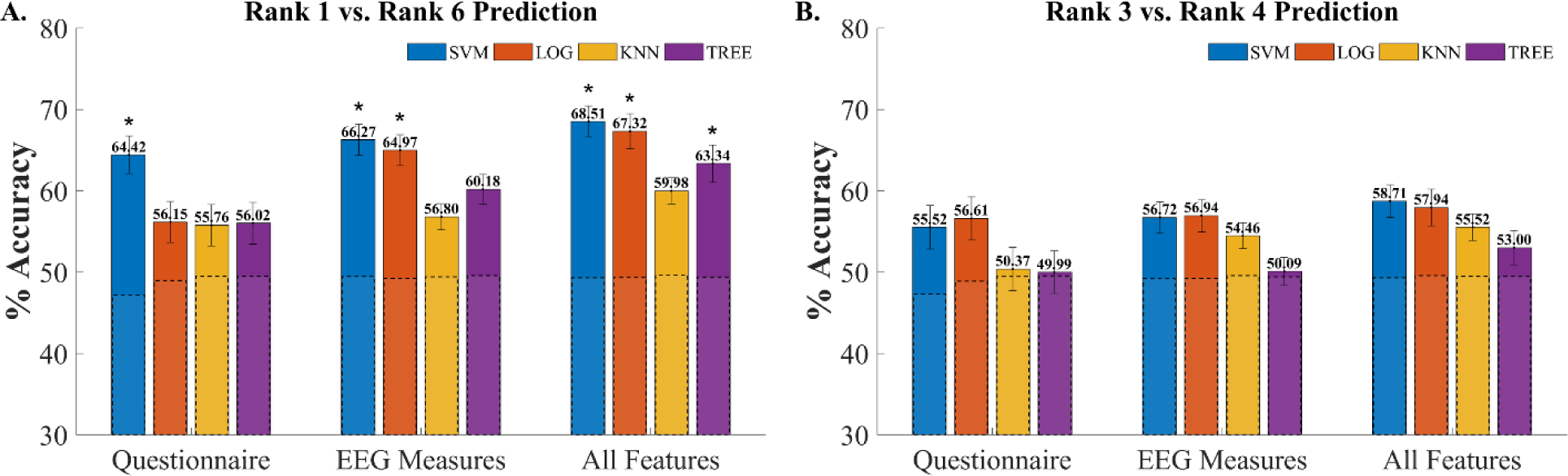
Binary Prediction Accuracies. Prediction of the most favorite products against the least favorite (**A**) yielded higher accuracies than when predicting between indistinct preferences (**B**). Also, accuracies were highest when EEG and Questionnaire features were combined, compared to each of them alone. Significance was obtained from permutation testing as explained in the methods (* = p < 0.05). Dashed bars indicate the mean of shuffled permutations accuracies. Error bars indicate Standard Deviation of the bootstrap distribution (Hesterberg, 2015).

Lastly, prediction accuracies did not differ from predictions on shuffled data when trying to predict between products ranked as 3 and 4. Hence, we inferred that neural measures were effective predictors only when products’ preferences were well disparate in subjects’ minds, such that they ranked them highest or lowest, rather than preferences they were more indifferent between. This is in concordance with the concept of “neural distance”, the idea that as the distance between preferences increase, the “neural distance” increases as well, and prediction accuracy gets better (I. Levy et al., 2011; Telpaz et al., 2015).

### Within-Subject Prediction – Multi-Class

We inspected the Root Mean Square Error (RMSE) results for perdicting all six subjects’ rankings, (**Figure 7.A**). This measure is calculated by summing the squared differences between each prediction to its corresponding true label, and then calculating the square root of the resultant sum. RMSE was used instead of prediction accuracy to account for the size of prediction errors between the six different rankings. That is, when the algorithm assigns a rank of 2 when the true rank is 3, this should be considered as a small error compared to when the algorithm assigns a rank of 6 when the true rank is 3. Note, that we lose this sensitiviy of the results when using simple prediction accuracies. Results for predicting the entire range of subjects’ rankings lead to similar conclusions as for the binary predictions; The combination of EEG and Questionnaire measures yeilds the lowest RMSE scores. This showed clearly that the neural measures assisted in prediction beyond the questionnaire on its own. Note, that the lowest RMSE scores were achived for ordinal models such as SVOREX and KDLOR which consider the ordinal rankings of the products (and not just their nominal label). Additionally, it seemed that the EEG measures performed slightly better than the questionnaire in this case.

**Figure 7.**
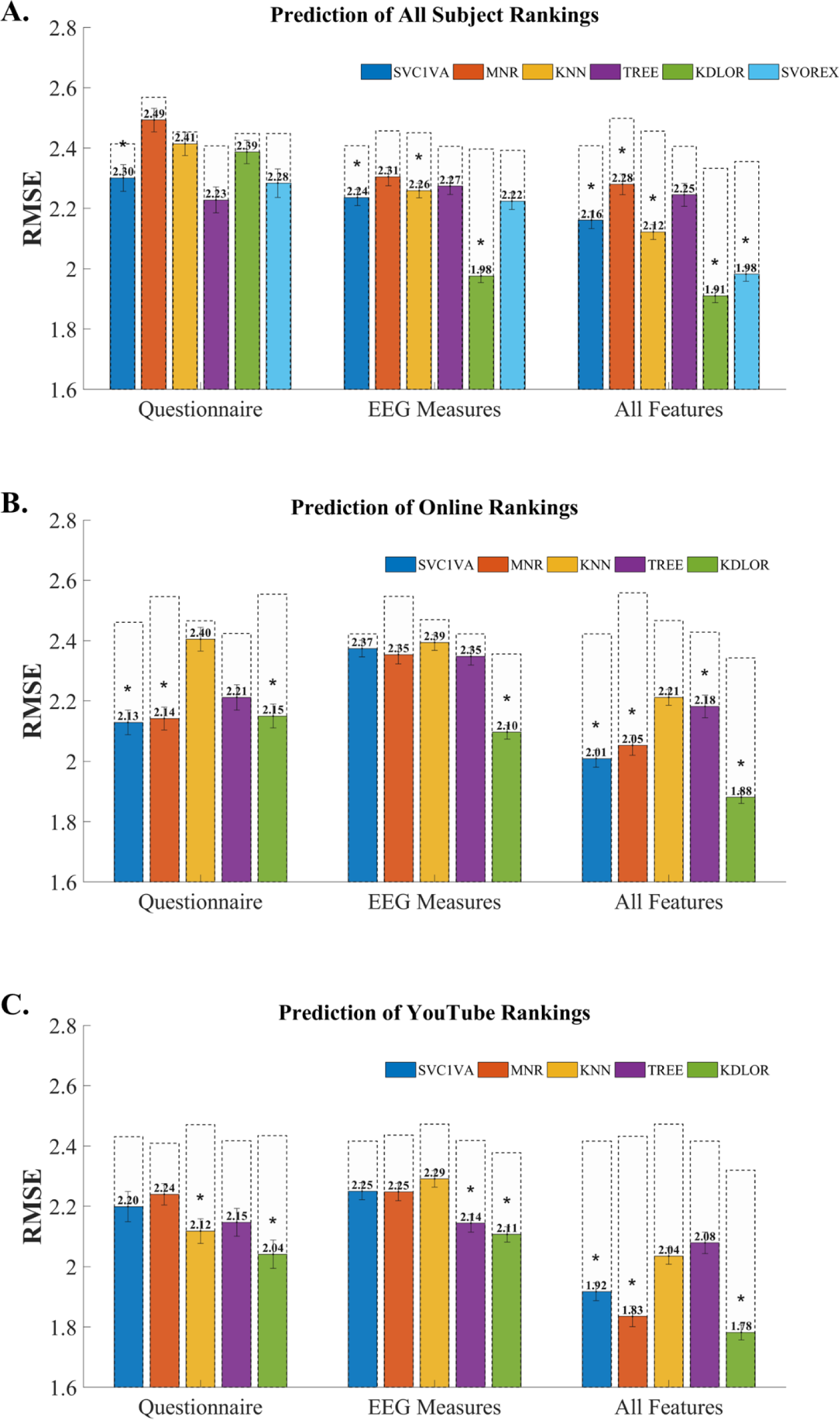
All Ranks Prediction RMSE. The graphs show RMSE scores for prediction of all six rankings when labels are (**A**) subjects’ rankings, (**B**) out-of-sample Questionnaire rankings, and (**C**) YouTube Metrics. Significance was obtained from permutation testing as explained in the methods (* = p < 0.05). Dashed bars indicate the mean of shuffled permutations accuracies. Error bars indicate Standard Deviation of the bootstrap distribution (Hesterberg, 2015).

### *Out-of-Sample Prediction – Multi-Class* – Online Questionnaire

Next, we attempted to predict the aggregate online questionnaire rankings, as a proxy for population preferences, based on the EEG and Questionnaire data from our subject pool. As this is what truely interests marketers and managers, we aimed to show that our EEG measures would again contribute to prediction of real market success of commercials, beyond traditional measures. Our results were that all models reached lower RMSE for the combined measures than with the questionnaire alone (**Figure 7.B**). This demonstrated that the EEG measures contributed to prediction of out-of-sample online questionnaire rankings, above and beyond the questionnaire.

### *Out-of-Sample Prediction – Multi-Class -* YouTube Metrics

To further strengthen the claim for the potential to predict market success from EEG measures, we also attempted to predict YouTube rankings of our product commercials, based on aggreagate YouTube metrics, as an additional proxy for population prefrences. When predicting YouTube rankings of product commercials (**Figure 7.C**), we obtained similar results as for the online questionnaire. The RMSE decreased when we combined the EEG with questionnaire data, compared to predicting based on the questionnaire alone.

In addition, it seems that the questionnaire measures are slightly better predictors than EEG measures, when each are used by themselves. Moreover, it is possible that the YouTube metrics are a better proxy for the population prefrences (as our subject sample pool are representing the population) than online questionnaires, as our models reached better prediction of YouTube rankings based on data from our subject pool. This could be due to the aforementioned biases and disadvantages of questionnaires, which may even affect an online cohort with a multitude of respondants. In contrast, YouTube metrics are more objective and reliable, their sample pool is much larger than can be obtained by online questionnaires, and in general are more ecological than generalizing based on individual preferences as obtained by online questionnaires. This could be of particular interest to marketers, as they often attempt to predict a campaign’s success by using self-report subjective online questionaires.

Importantly, in both the YouTube and Online Questionnaire predictions, we found that the best model (which yeiled the lowest RMSE between all predictions) was KDLOR. This is probably because this model is the most acute, as it is the only “Ordinal-type” model, which take into accout the rank order of the labels for training and predicting. The other ordinal mode, “SVOREX”, was not used due to technichal difficulty (crashes) when attempting to apply it. Lastly, it is worth noticing that EEG measures performed better by themselves than the questionnaire measures when predicting the subjects’ rankings, while they performed sligthly worse when predicting Online and Youtube rankings. This could mean that the EEG measures are more tightly linked to individual prefrences than aggregate choices.

## Discussion

In the current study, we show that using a combination of various features from several EEG signal types and advanced machine learning models increase the predictive power of subjects’ future choices above and beyond a traditional marketing questionnaire. Moreover, this is the first study to demonstrate that we can use EEG signals to predict rank ordered preferences of an out-of-sample cohort of subjects and the overall popularity at the population level of the commercials. Lastly, we highlight several important prediction challenges when using a machine learning approach that if not accounted for will inflate prediction accuracies, cause overfitting and could lead to biased and inaccurate conclusions regarding the prediction outcomes (as illustrated in **Figure 2.D**).

For some of our prediction attempts, we were able to show significant prediction accuracies when we used the combination of several EEG measures by themselves. We showed a similar result when using only the features from the questionnaire. However, when we wanted to predict subjects’ future rank ordered preferences, it was the composite of both EEG and questionnaire features that had the highest significant predictive accuracy. These results suggest that each type of feature captures a different aspect of the valuation process and the combination gives rise to a better prediction. Hence, we emphasize that neuroscientific tools should be combined with standard measures for better prediction.

On the other hand, in our data, we showed that the EEG measures alone were better in predicting subject’s individual rankings, be it all six rankings or just two rankings, than the questionnaire alone. Meanwhile, the questionnaire alone outperformed the EEG measures for population metrics predictions. A possible explanation for this is that neural measures of different subjects were more indicative of their own personal preference than the marketing questionnaire they filled, while the questionnaire on its own is somewhat more predictive of a similar online questionnaire and YouTube metrics. This result could suggest that neural measures contain hidden information regarding general preferability in individual subjects, which questionnaire responses might not capture. Questionnaires may be more rigid and biased, and contain the disadvantages mentioned in the introduction. However, it should be noted that our EEG measures included 15 features, while from the questionnaire we extracted 3 features only, which were the strongest predictors we were able to extract as to not give our neural measures an unfair advantage. It is possible that further scrutiny, exploration and optimization of the questionnaire responses would yield varying results. Yet, this is exactly part of our critique of questionnaires. Ideally, there could exist a perfect questionnaire which captures individual and population preferences precisely in a particular marketing setting, but it may take an endless amount of time and effort calibrating such questions for each setting, while the literature is already beginning to converge on powerful, general and objective neural predictors of preference.

In contrast to findings from previous experiments (Venkatraman et al., 2015), we show that EEG measures contributed substantially to prediction. This was due to three main innovations explored in this paper. The first was the use of multiple complex EEG features, which gave prediction models broader information regarding subjects’ neural activity and improved their ability to capture subjects’ valuations of product commercials. The second novel aspect was searching beyond standard regression techniques, into novel prediction models from the immensely promising world of machine learning. This allowed us to attempt various modelling approaches which could better suit our prediction task. Lastly, we accounted for possible errors in the analysis process that cleaned our predictions and strengthened their interpretation.

Applying machine learning approaches on EEG data can lead to several caveats, which we aimed to identify and address. EEG activity can differ widely between subjects and trials, requiring appropriate normalization within and between subjects for these models to be used correctly. The multi-leveled nature of experimental datasets also demanded careful handling to avoid leakage, as did any transformations on the data which could be considered peaking and possibly yield falsely inflated prediction rates (see **Figure 2** and methods for details).

Importantly, our modeling approach allowed us to avoid the issues of multiple comparisons, by relying on previous literature for feature creation while utilizing all measures for all frequency bands in our predictions. Moreover, we examined ecological and market-relevant stimuli, without the need to subjectively judge commercials or specific scenes within them. This was done by averaging over powers during the entire commercial (FBP, Asymmetry), or preforming analysis that is based on the entire commercial timeline (ISC).

Another significant conclusion from our inspection of EEG strength as a predictor of preferences, was the influence of between subject variation in preferences on performing meaningful predictions. For within-subject prediction, if subjects were highly correlated with one another, such that most like and dislike the same products, then predictions become closer to predicting the product commercial identity rather than subject-specific ranking of that commercial. That is, the EEG measures could be reflective of the commercials objective characteristic, such as its length, saliency, brightness, motility and others, rather than individual preferences, and there would be no way to tell apart the two possible interpretations of the EEG measures’ predictability.

For out-of-sample predictions it is even harder to tell them apart. First, it is easier to predict an overall population product ranking than subject specific choice. When attempting to predict subject-specific preferences, the rank of a product can change for every subject. On the other hand, when attempting to predict population-level metrics the rank of a product remains the same across all subjects – the averaged rank allotted to the product by the population. Then, if subjects’ rankings correlate well with the population rankings, i.e. everyone likes and dislikes the same products, the EEG measures only need to capture a pattern of response to the objective attributes of the product commercials, rather than any value-related signals, in order to predict population ranks successfully. This interpretation is particularly strong for small sets of products, but its relevance decreases the more products are included in the prediction attempt. For six products, our model could be predicting the population ranking simply as an identifier of the general cortical activity in response to each commercial, rather than an ordinal value. For 50 products, it would be much harder to say that population rankings are simply identifiers, as their ordering becomes a much more significant element of prediction. If neural measures continue to predict population rankings with an increasing number of products, it can no longer be said that the identity and characteristics of products are being used to predict, but rather their order of value. Therefore, future studies should focus on strengthening supporting evidence for powerful neural predictors using larger quantities of stimuli.

Once neural predictors of value are adequately established, then they can be utilized on real-world problems that interest marketers and managers, which usually involve only a small set of closely similar stimuli.

This issue presents a limitation on predicting population commercial success in general, but it is not an issue particular to EEG, but to anyone attempting to predict population commercial success from a small sampled group. Our study showed the contribution that EEG has for out-of-sample prediction that could be easily replicated for any marketing decision – be it choosing out of different versions of a certain commercial, differing brands of a product type, between varying product categories, or otherwise.

Moreover, the research community could benefit mainly from our results regarding within-sample predictions. These showed that substantial success can be achieved by using multiple complex measures of EEG recordings that are based on previous literature. This suggests that the neural correlates of value, as measured by EEG, are probably a combination of various measures, which reflect different cognitive aspects of the valuation process, and are not restricted to a specific unique signal, such as the BOLD signal measured in fMRI (Bartra et al., 2013; D. J. Levy & Glimcher, 2012). Researchers should also be incentivized to discover novel ways to process the EEG signal and develop new measures to capture aspects of value information independent of existing measures, to further enhance EEG-based prediction.

In conclusion, although considerable knowledge regarding consumer preferences is attainable through traditional marketing techniques, such as questionnaires and focus groups, many marketing strategies still prove relatively unsuccessful in aiding managers and to make business and marketing decisions (Hamel & Prahalad, 1994; Martin, 1995; Ovans, 1998). The relevant neuroscientific literature has found numerus evidence that link the neural measures we used to consumer preferences. Moreover, we apply novel data science methodology and techniques that align with industry standard prediction modeling. Hence, our research expands the manager’s toolbox, with a cost-effective and practical tool, EEG, with which they can access customers’ cortical activity in order to execute managerial decisions empirically, possibly better than what traditional behavioral measures could provide. Practitioners can utilize our findings to predict responses to new company products, to modifications in existing products, to specific tactics or to various marketing campaign options and branding decisions, before substantial investment in media spending. By achieving accurate marketing performance predictions, as our results with population prediction imply, managers could drastically decrease failures or uncertainties in their strategy, increase their marketing effectiveness, broaden their audience, improve brand image, and maximize their return on investment.

## Acknowledgements

We want to thank all the Dino Levy’s lab members for their help and guidance in many discussions. We also thank Yoav Zeevi, a Ph.D. student of Professor Yoav Benjamini, for his guidance with statistical tests performed throughout the paper.

## Author contributions statement

AH performed the data analysis, created and optimized the prediction models and modelling challenges. TS and MSS designed and conducted the experimental procedure. SK conducted the experiment also, performed EEG preprocessing and manual artifact rejection. AH wrote the manuscript with the aid and supervision of DL. All authors read and approved the final manuscript.

## Supplementary information

- All Prediction Results.
- Marketing Questionnaire

## Additional information Competing financial interests

